# The temperature-dependent predatory strike of Odonata larvae

**DOI:** 10.1101/2020.12.08.416289

**Authors:** Alexander Koehnsen, Hannah-Lena Tröger, Stanislav N Gorb, Sebastian Büsse

## Abstract

The larvae of Odonata are limnic predators capable of catching their prey using a highly modified mouthpart – the labium. Driven by a unique dual catapult mechanism, the apparatus can reach peak accelerations of up to 114.5m/s^2^. Yet little is known about the kinematics of the predatory strike in an ecological context. Here we show how different ambient temperatures affect the predatory strike and the avoidance reaction of prey items of Odonata larvae. We found that the extension velocity of the labial mask decreases significantly with the ambient temperature both in dragonflies and damselflies. However, temperature has lesser impact on the predatory strike itself than on directly muscle driven movements in both the predator and prey items. This contradicts the previous assumption that catapult mechanisms in insects are unaffected by temperature. Our results indicate that the prehensile labial mask is driven by a series-elastic catapult; a mechanism similar to the temperature dependent jump of frogs, where muscle and spring action are tightly linked. Our study provides novel insights into the predatory strike of Odonata larvae and offers a new ecological perspective on catapult mechanisms in arthropods in general.

## Introduction

Odonata (dragonflies and damselflies) are remarkable predators. While the imagines roam the air – catching prey on the wing – while the larvae catch aquatic prey in their benthic habitats (Corbet, 1999). These larval predators rely on a unique mechanism for prey capture, using a highly modified mouthpart – the labium (Snodgrass, 1955; Büsse et al., 2017), capable of rapid extension (Pritchard, 1965; Büsse et al., 2020). According to Pritchard (1965), the predatory strike consists of four distinguishable phases (Figure 1): 1) the labial palps open in preparation for the predatory strike; 2) the labium extends; 3) labial palps close around the prey item; 4) the labium retracts. The driving force of the prehensile labial mask consists of two synchronised catapults (Büsse et al., 2020). This specialised apparatus enables the larva to catch prey up to its own body size (Corbet, 1999). Tadpoles or even small fish are not an uncommon prey for large larvae like hawkers (Aeshnidae) (Pritchard, 1964; Jara and Perotti, 2010). Small water bodies represent a highly challenging habitat, as temperature fluctuates drastically over the course of a year (Wiggins et al., 1980) and even diurnally (Corbet, 1999). As the activity of poikilothermic animals depends on the ambient temperature (Huey and Kingsolver, 1989), fluctuating ambient temperatures represents a challenge for successful prey capture, as the predator efficiency changes with temperature (Kruse et al., 2008). Studies in other arthropods have shown that ingestion efficiency (food uptake divided by metabolic rate) decreases with higher temperatures, because food uptake remains constant but the metabolic rate increases (Rall et al., 2009; Vucic-Pestic et al., 2011). This seems to be a general phenomenon in poikilothermic animals (Rall et al., 2012). Therefore, effective prey capture at lower temperatures might be beneficial for energy uptake and growth. However, the muscle contraction time simultaneously increases at lower temperatures, considerably reducing the speed of muscle driven movements in poikilothermic animals (Huey and Kingsolver, 1989; Josephson, 1993) and some models suggest that prey is at an advantage when temperatures are low and both predator and prey are ectothermic (Dell et al., 2014).

**Figure 1:**
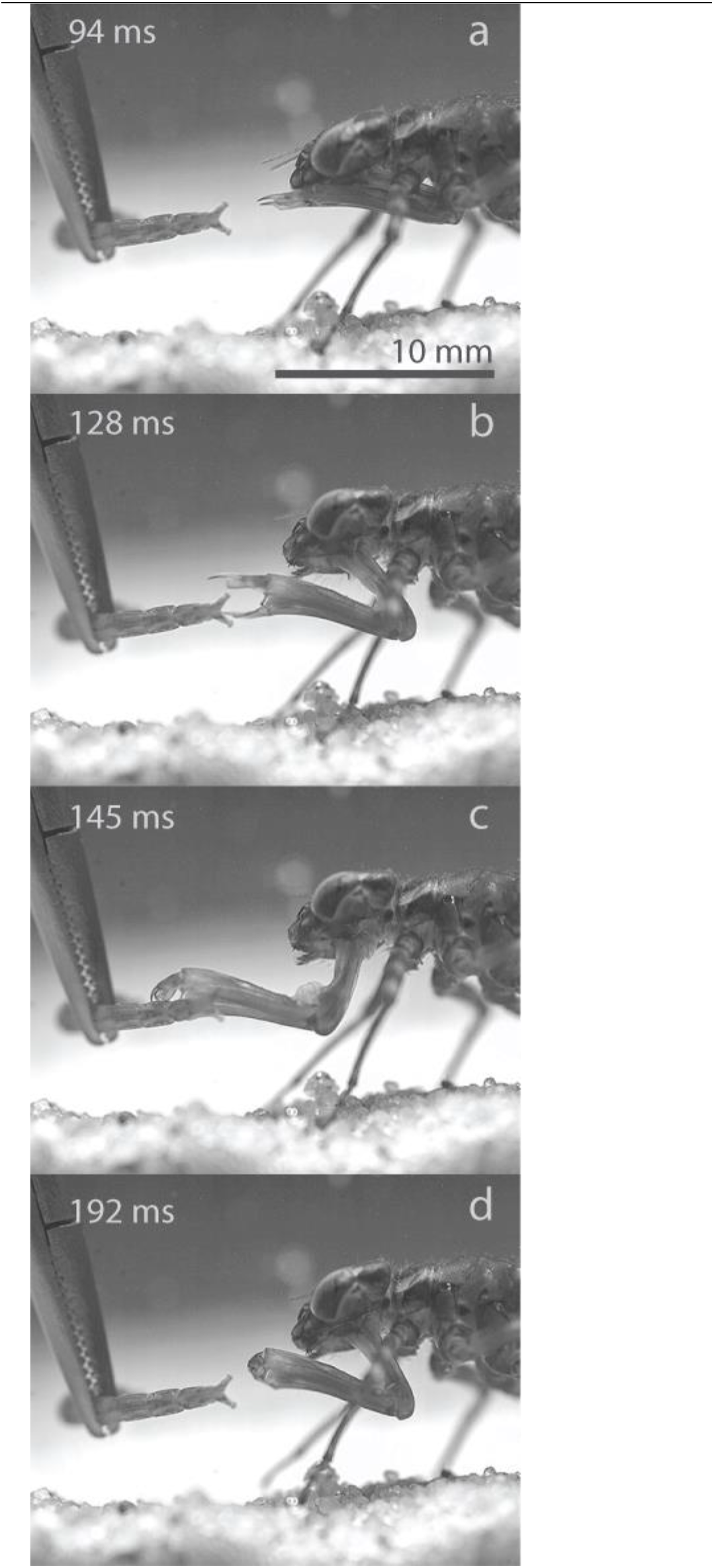
Overview of strike phases. Exemplary strike of *Anax imperator.* a) Phase I, the opening of the labial palps. 94ms after opening started. b) Phase II, the projection of the prehensile labial mask. c) Phase III, the closure of labial palps around the prey item (in this case they miss). d) Phase IV, the retraction of the prehensile labial mask. Extracted frames from high-speed footage captured at 5400 fps.

The indirectly muscle driven catapult system of the prehensile labial mask (Büsse et al., 2020) might hold a key advantage under the described circumstances. Catapult mechanisms are not directly powered by a muscle, but rather preloaded and energy is stored using an elastic structure (Gronenberg, 1996; Ilton et al., 2018). In the case of Odonata larvae, two sclerites inside the prehensile labial mask serve as springs and the surrounding cuticle might be involved in energy storage as well (Büsse et al., 2020). The contraction time of muscles should therefore not influence the velocity of a catapult driven movement (Gronenberg, 1996). An earlier study has already shown that the odonate predatory strike is affected by temperature changes (Quenta Herrera et al., 2018), yet the authors focused on the ecological implications of this effect. In this study, we try to find a biomechanical explanation for the temperature dependence and compare the performance loss at lower temperature to common prey items.

In this study, we assess the kinematics of the predatory strike of Odonata larvae at two different temperature ranges (low: 9.5-13.5°C high: 19-25°C) using high-speed videography. We used specimens of two Anisoptera species *Anax imperator* LEACH, 1815 and *Sympetrum* sp. NEWMAN, 1833 as well as one Zygoptera species *Erythromma najas* (HANSEMANN, 1823), allowing for a comparison of strike characteristics between these groups. We aimed to investigate whether the strike kinematics between Anisoptera and Zygoptera differ, as well as how the predatory strike is affected by temperature. Finally, how does the decrease in the strike performance that was demonstrated by Quenta Herrera et al., (2018) affect the predatory strike in comparison to the escape reaction of prey items? For this purpose, we analysed the avoidance reaction and swimming speed of typical prey items of Odonata larvae at the same two temperature ranges.

Answering these questions will improve our understanding of the role of this unique prey capturing mechanism in the success of Odonata larvae as key predators in their habitats. This will also provide information on the biomechanics of the predatory strike and their ecological implications within the predator-prey relationship. Finally, we will offer a new perspective on the functional morphology of the catapult mechanisms in arthropods by providing some insights on why catapults in arthropods may be temperature dependent.

## Materials and methods

### Specimens

Larvae of *Anax imperator* (Anisoptera: Aeshnidae) were collected in aquaculture ponds in Oeversee, Schleswig-Holstein, Germany. Nine specimens of *Sympetrum* sp. were collected in a small pond close to Knoop, Schleswig-Holstein, Germany. 13 Specimens of *Erythromma najas* (Zygoptera: Coenagrionidae) were collected in a pond in the Botanical Garden in Kiel, Schleswig-Holstein, Germany. All specimens were collected with the permission of the ‘Landesamt für Landwirtschaft, Umwelt und ländliche Räume Schleswig-Holstein‘. Specimens were kept in individual aquariums at 8°C to prevent premature emergence and fed with chironomid larvae (Diptera: Chironomidae). Before experiments, the Odonata larvae were slowly acclimatised to the temperature range of the experiment to avoid temperature shocks and measuring errors. For all specimens, head length and width were determined to provide dimensions for scaling.

### High-speed videography

High-speed video recordings of the predatory strike were performed in a small aquarium (210×100×105mm). Ten specimens of *Anax imperator* (Odonata: Anisoptera), nine specimens of *Sympetrum* sp. (Odonata: Anisoptera) and 13 specimens of *Erythromma najas* (Odonata: Zygoptera) were used. Chironomid larvae were manually presented as prey items. Experiments were performed at two temperature ranges: 9.5-13.5°C and 19-25°C. The water temperature was constantly monitored during the experiments using an analogous thermometer. We recorded strikes of four specimens of *A. imperator* at the high-temperature range, and of six specimens at the low-temperature range. Nine specimens of *E. najas* were used at the high-temperature range, and four specimens of *E. najas* were used at the low-temperature range. In total, 80 high-speed recordings were made with a Photron Fastcam SA1.1 (model 675K-M1; Photron, Pfullingen, Germany) equipped with a 105 mm/1:2.8 macro lens (Sigma, Tokyo, Japan) mounted on a Manfrotto-055 tripod with Manfrotto-410 geared head (Manfrotto, Spa, Italy). Illumination was provided using two Dedocol COOLT3 halogen spotlights (Dedotech, Berikon, Switzerland). The footage was captured with a resolution of 1024×1024 pixels at 5400 frames per second and saved as a 16-Bit TIFF image-stack. To obtain average velocity and acceleration, the distance travelled by the tip of the prementum from the start of the extension phase until the labial palps start to close was determined using the measurement tool of Adobe Photoshop CS6 (Adobe Systems Software, San José, CA, USA).

### Experiments on temperature-dependent prey behaviour

Two typical prey items of Odonata larvae were collected in a pond in the Botanical Garden in Kiel, Schleswig-Holstein, Germany. Larvae of Ephemeroptera (Insecta: Ephemeroptera) and larvae of *Erythromma najas* (Odonata: Zygoptera) of early instars were used. The swimming velocity was determined using high-speed videography with the previously mentioned setup, but recordings were made at 1000 fps. An avoidance reaction was triggered by a mechanical stimulus. To obtain average velocity and acceleration, the tip of the centre of mass was tracked according to Koehnsen et al. (2020).

### Statistical analysis

A Mann-Whitney Rank Sum test was used to compare the changes in average strike velocity as sample sizes were not equal and data were not normally distributed (criterion for normal distribution: Shapiro Wilk test, p > 0.05). For the comparison of average swimming speeds in prey items, a Welch t-test was used, as sample sizes were equal and data were normally distributed. All statistical analyses were performed in R Studio Version 3.3.1 (The R Foundation for Statistical Computing, Vienna, Austria). N indicates the number of biological replicates (specimens).

### Estimation of drag force

To estimate the effect of temperature change on the occurring drag force *F_d_*, we calculated the influence of the density and drag coefficient. The calculations are shown in supplementary material S1. Physical parameters of water (density and viscosity at different temperatures) were taken from Rumble et al. (2017).

## Results

We performed feeding experiments at a constant temperature range (19°-25°C) with specimens of *A. imperator* (Odonata: Anisoptera) and *E. najas* (Odonata: Zygoptera) using chironomid larvae as prey items and analysed the predatory strike using high-speed videography. The average distance travelled by the prementum during protraction is longer in *A. imperator* (4.7 +−1.3mm) than in *E. najas* (1.9 +−0.03mm). From the distance travelled by the prementum and the duration of the protraction, average velocity and average acceleration were determined (Figure 2). While there is no significant difference in velocity (t-Test, p=0.13, N=17), the acceleration of the prementum is significantly higher in *E. najas* (Mann-Whitney Rank Sum Test, p<0.001, N=17).

**Figure 2:**
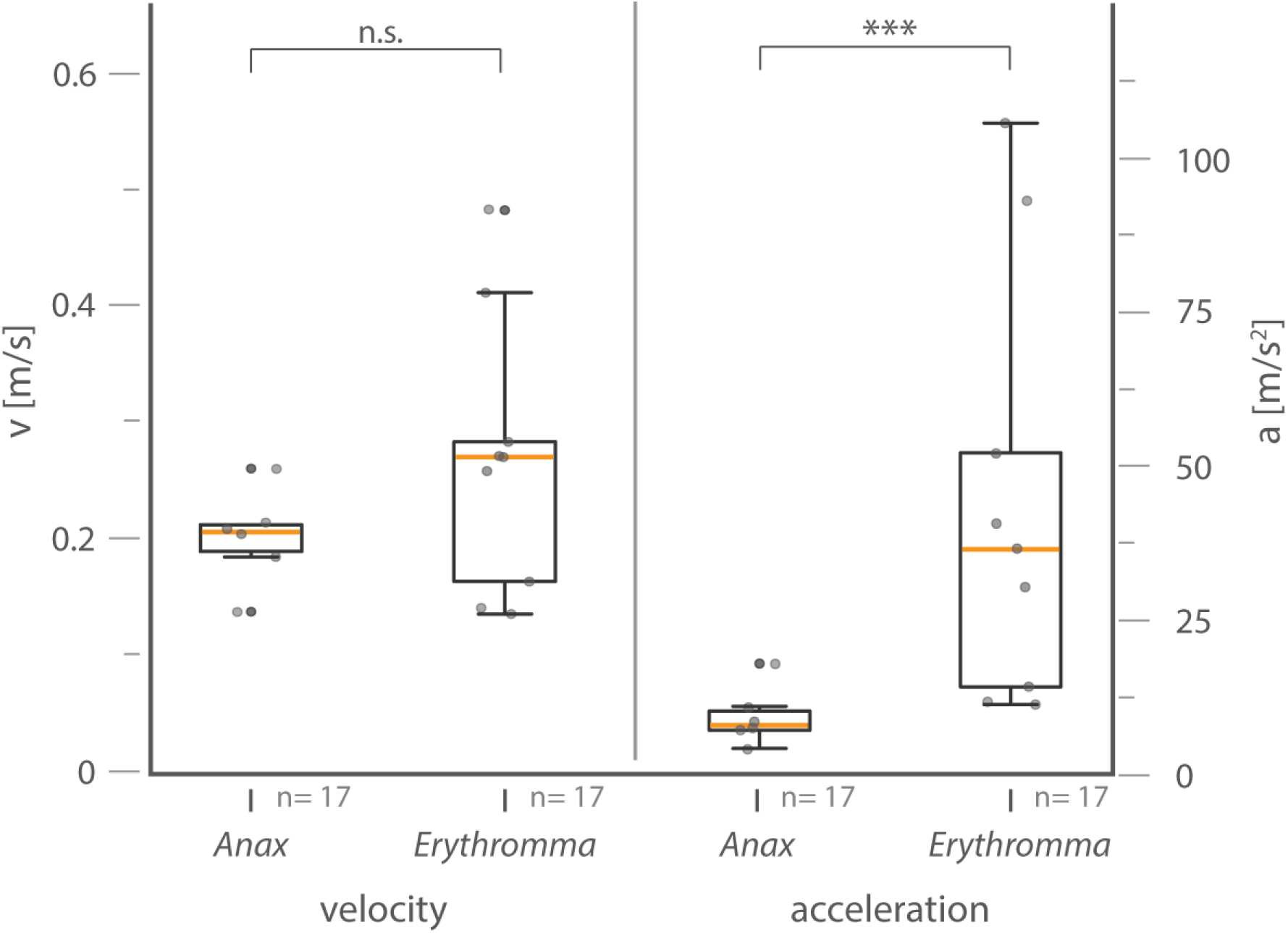
Differences in strike velocity and acceleration between *Anax* and *Erythromma*. Velocity: t-Test, p=0.13, N=17; acceleration: Mann-Whitney Rank Sum Test, p<0.001, N=17. While the median velocity is comparable in both groups (0.21m/s in *A. imperator* and 0.28m/s in *E. najas*), the median acceleration is around four times higher in *E. najas* (9.4 m/s^2^ in *A. imperator* and 39.9m/s^2^ *in E. najas*). Box ends define 25^th^ and 75^th^ percentiles, with a line highlighting the median and error bars at the 10^th^ and 90^th^ percentiles. Dots indicate individual data points.

We recorded predatory strikes of *A. imperator*, *Sympetrum* sp. and *E. najas* at two temperature ranges (low-temperature range 9.5-13.5°C, high-temperature range 19-25°C). The results show that the velocity of the predatory strike is dependent on temperature (Figure 3). Strike velocity decreased in all three cases significantly (*Sympetrum sp.* Mann-Whitney Rank Sum Test, p=0.004 N=9 *A. imperator*: Mann-Whitney Rank Sum Test, p=0.006 N=17 (cold), N=16 (warm), *E. najas*: t-test, p<0.001 N=17 (cold), /N=12 (warm)) with decreasing temperature. We found that the average protraction velocity differs between the two temperature ranges by 34% in *A. imperator,* 34% in *Sympetrum* sp., and 61% in *E. najas*.

**Figure 3:**
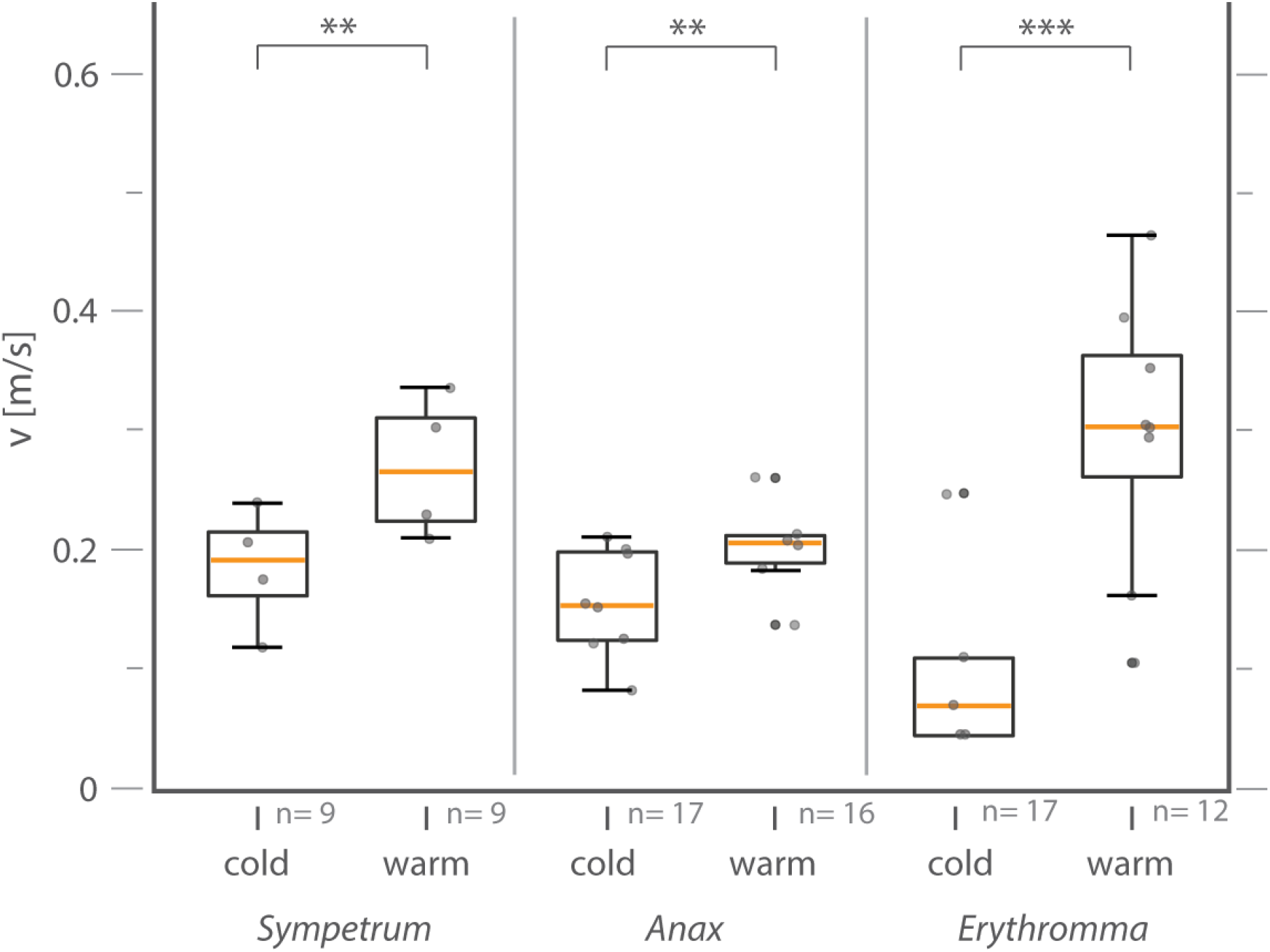
Strike velocity decreases significantly with temperature. Comparison of the average velocity of the prehensile labial mask during extension (phase 2) in three Odonata larvae at two temperature ranges (low-temperature range (9.5-13.5°C); high-temperature range (19-25°C)). In all three species, strike velocity decreases significantly with temperature (*Sympetrum* sp.: Mann-Whitney Rank Sum Test, p=0.004 N=9, *A. imperator*: Mann-Whitney Rank Sum Test, p=0.006 N=17/16, *E. najas*: t-test, p<0.001 N=17/12). Box ends define 25^th^ and 75^th^ percentiles, with a line highlighting the median and error bars at the 10^th^ and 90^th^ percentiles. Dots indicate individual data points.

The duration of individual strike phases differs at the two ambient temperature ranges as well (supplementary figure 2). Both the duration of phase 1, during which the labial palps open and the duration of phase 2+3, during which the prehensile labial mask protracts, increase significantly (*A. imperator*: phase 1: Mann-Whitney Rank Sum Test, p<0.001 N=17 (cold), N=16 (warm); phase 2: t-test, p<0.001 N=17 (cold), N=16 (warm); *E. najas*: phase 1: Mann-Whitney Rank Sum Test, p<0.001 N=17 (cold), N=12 (warm); phase 2: Mann-Whitney Rank Sum Test, p<0.001 n=17 (cold), N=12(warm)). When the relative increase in average phase duration is compared, phase 1 increases in *A. imperator* by 74.0ms (50%), while the duration of the protraction only increases by 15.1ms (30%). The same trend was observed in *E. najas*, where the average duration of the phase 1 increases by 31.1 ms (71%), while the duration of the phase 2 increases by 20.9 ms (61%).

To compare the performance of the catapult driven prehensile labial mask to the escape behaviour of various prey items, we recorded footage from Zygoptera and Ephemeroptera larvae, typical prey items of Odonata larvae. We did not use Chironomid larvae, as they did not exhibit a quantifiable escape behaviour. 15 clips were recorded at each temperature range (19-25°C, 9.5-13.5°C), from which the average swimming velocity was determined (Fig. 4). In both cases, the swimming velocity decreased significantly between the warm and cold temperature range (Zygoptera larvae: t-test, p<0.001, N=15; Ephemeroptera larvae: t-test, p<0.001, N=15). The relative decrease in swimming velocity due to temperature decrease differs between the two taxa: Ephemeroptera larvae show a decrease by 79%, while Zygoptera larvae show a decrease by 45%.

**Figure 4:**
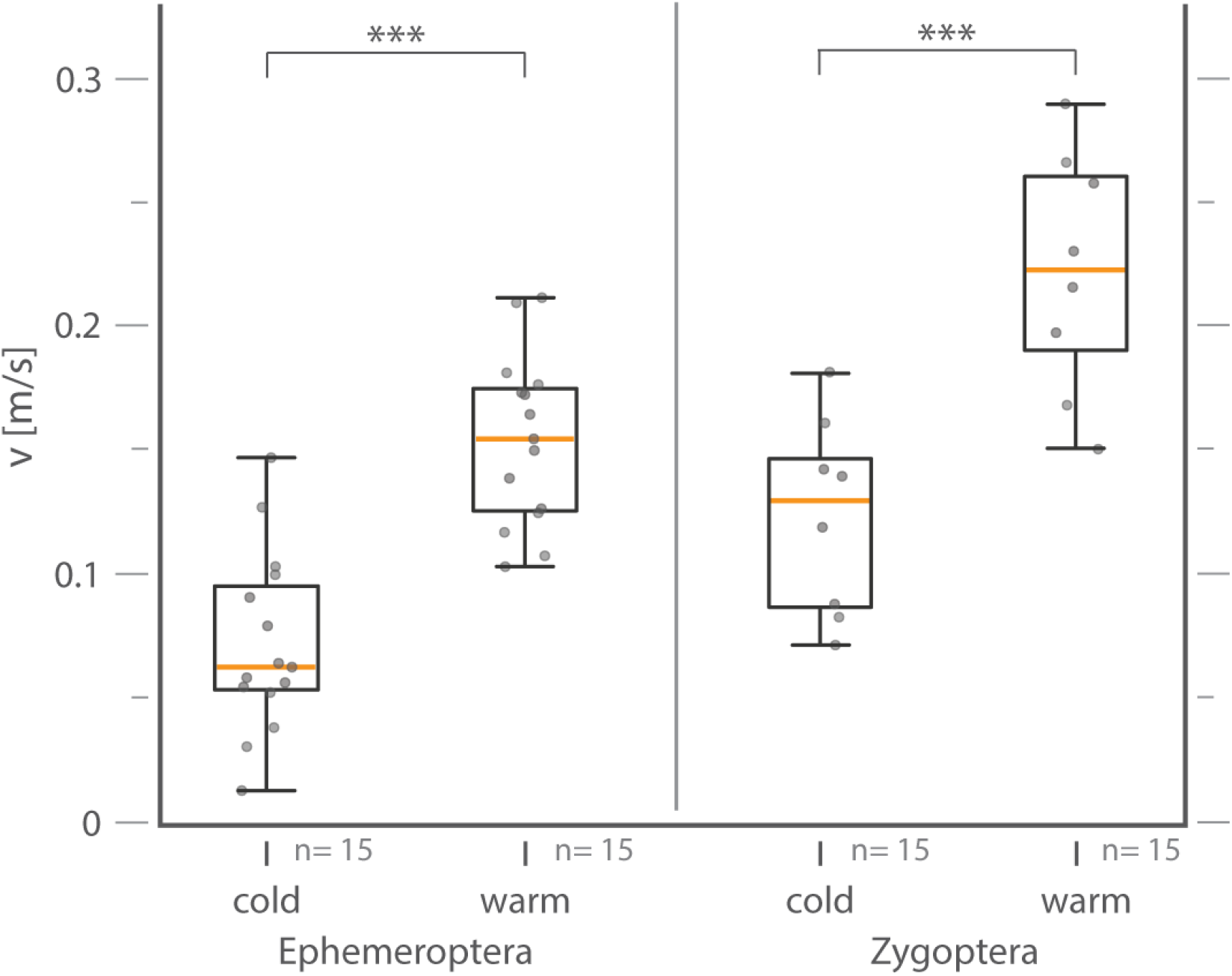
Average swimming velocity of potential prey items at different ambient temperatures: low (9.5-13.5°C) and high (19-25°C). Data was obtained using high-speed videography. In both cases, the average swimming velocity decreased significantly with a decrease in ambient temperature (Zygoptera larvae: t-Test, p<0.001, n=15; Ephemeroptera larvae: t-Test, p<0.001, n=15). Box ends define 25^th^ and 75^th^ percentiles, with a line highlighting the median and error bars at the 10^th^ and 90^th^ percentiles. Dots indicate individual data points.

According to the drag equation, temperature-depending variables affecting drag forces are (1) the water density *ρ* and (2) the drag coefficient *C*_*d*_ due to change of the Reynolds number (*Re*). Temperature decrease from 20°C to 10°C leads to the *ρ* change of −0.001495 g/ml, which is less than 0.15%. As *ρ* is incorporated into the drag equation on a linear basis, a change in temperature to 10°C would, if all other variables remain equal, change the drag around the prehensile labial mask by less than one percent (supplementary material S1). The Reynolds number decreases by around 23% in *Erythromma* (~ 650 - ~ 500) and *Anax* (~ 2200 - ~ 1700), when temperature decreases by 10°C (supplementary material S1).

## Discussion

We analysed the strike performance of the anisopteran species *Anax imperator, Sympetrum* sp. and the zygopteran species *Erythromma najas* by assessing the maximum strike velocity and maximum acceleration. Our results have shown that although there is no significant difference in strike velocity between species, the acceleration of the prehensile labial mask is significantly higher in *E. najas* (Fig 2). *E. najas* is a much smaller species, and therefore also the mass of the prehensile labial mask is considerably lower than in *A. imperator* and *Sympetrum* sp. Lower mass and corresponding lower inertia allow for higher accelerations. This trend consistently found in high-speed movements of animals: small yet fast movements are combined with extreme accelerations (Gronenberg, 1996; Ilton et al., 2018). It is necessary for structures involved in these movements to provide higher accelerations, as the distance that can be used to accelerate decreases with decreasing size of the animal. The results of our experiment reflect this trend very well, as all three species have about the same strike velocity, but the smaller prehensile labial mask of *E. najas* requires significantly higher accelerations to achieve the same velocity within the shorter strike distance.

We were able to show that the strike velocity decreased significantly with temperature in all three species studied (Fig 3). The average protraction velocity decreased by 34% in both *A. imperator* and *Sympetrum* sp. and by 61% in *E. najas.* This result is consistent with previous study on dragonfly larvae (Quenta Herrera et al., 2018) and contradicts the previous assumption that the predatory strike is not affected by temperature (Tanaka and Hisada, 1980). This result is counterintuitive, because catapult driven mechanisms in arthropods are generally not assumed to be temperature-dependent as energy release from springs should be independent from muscle contraction time (Gronenberg, 1996). In poikilothermic animals, the contraction time of a muscle increases with a decrease in temperature, wherefore the activity of poikilothermic animals decreases as well (Huey and Kingsolver, 1989; Josephson, 1993). Catapult mechanisms are powered by musculature, yet not directly. An elastic structure is preloaded by muscle power and acts as a storage unit for potential energy until the energy is released by a trigger system (Gronenberg, 1996; Ilton et al., 2018). Therefore the contraction time of a muscle should not influence the velocity of a catapult driven system (Gronenberg, 1996).

Hence different explanations have been brought up, such as the influence of increased drag due to higher fluid viscosity at a lower temperature has been discussed previously as a hypothesis for the temperature-dependent performance decrease (Quenta Herrera et al., 2018). We estimated the impact of temperature changes using the drag equation (for details on the drag equation, we recommend Hoerner, 1965; Vogel, 1997). The difference in drag force amounts to less than 1%, which does not suffice to explain the difference in strike velocity. The Reynolds number is likely to change due to temperature-dependent viscosity changes as well, which might influence the drag coefficient C_d_. If our results are compared with literature data of C_d_ changes of a sphere at different Reynolds numbers (Hoerner, 1965), C_d_ remains relatively constant at the range of values for *Anax*, but tends to change between around 0.2 and 0.4 for the *Re* of *Erythromma* which might explain the greater decrease in strike velocity of this species, as observed in our experiments (Fig. 3). In conclusion: Although being only a very rough estimate, the results suggest that these changes might play a minor role at best, but are unlikely to be sufficient to explain the distinct decrease in velocity. Further research beyond the scope of this study would be necessary to quantify the changes of flow dynamics at these different temperatures.

We hypothesise that the temperature dependence can be explained by a stronger interaction between spring and musculature: series elastic power modulation(Galantis and Woledge, 2003; Yesilevskiy et al., 2015, Haldane et al., 2016).Catapult mechanisms can be divided into two basic categories: parallel elastic and series elastic mechanisms (Galantis and Woledge, 2003; Yesilevskiy et al., 2015). A schematic of both principles after Yesilevskiy et al. (2015) is shown in figure 5. A parallel elastic mechanism is based on a spring that is preloaded by a motor (muscle) for a longer period of time. For the launch, the motor is decoupled and the catapult is only projected by the energy stored in the spring (Kovač et al., 2010; Yesilevskiy et al., 2015; Haldane et al., 2016). Such a mechanism would be completely independent of muscle contraction times and can be found for example in fleas (Sutton and Burrows, 2011). Serial elastic catapults are slightly different in function. Here muscle contraction can be subdivided into two phases: initially, the energy transferred to the moving part is lower than the energy invested by the muscle. The difference is stored by deforming an elastic structure. In the second phase of the motion, the energy is returned as the elastic structure recovers, raising the peak power output above the power output of the muscle (Galantis and Woledge, 2003; Yesilevskiy et al., 2015; Haldane et al., 2016). Such a series elastic mechanism can reach its peak power after the action of the motor is completed as shown via a robotic model by Haldane et al., (2016). This pattern with no muscle activity during parts of the motion matches the observations of Tanaka & Hisada (1980) on the predatory strike. The tight link of muscle and spring activity in a series elastic system might explain the performance deterioration caused by lower temperature. A similar effect has been shown for vertebrates, where the muscle tendon complex of the frog *Rana pipiens* SCHREBER, 1782 resembles a series elastic catapult mechanism (Astley and Roberts, 2012). Yet it has been shown that its jumping performance depends on temperature (Renaud and Stevens, 1983; Hirano and Rome, 1984).

**Figure 5:**
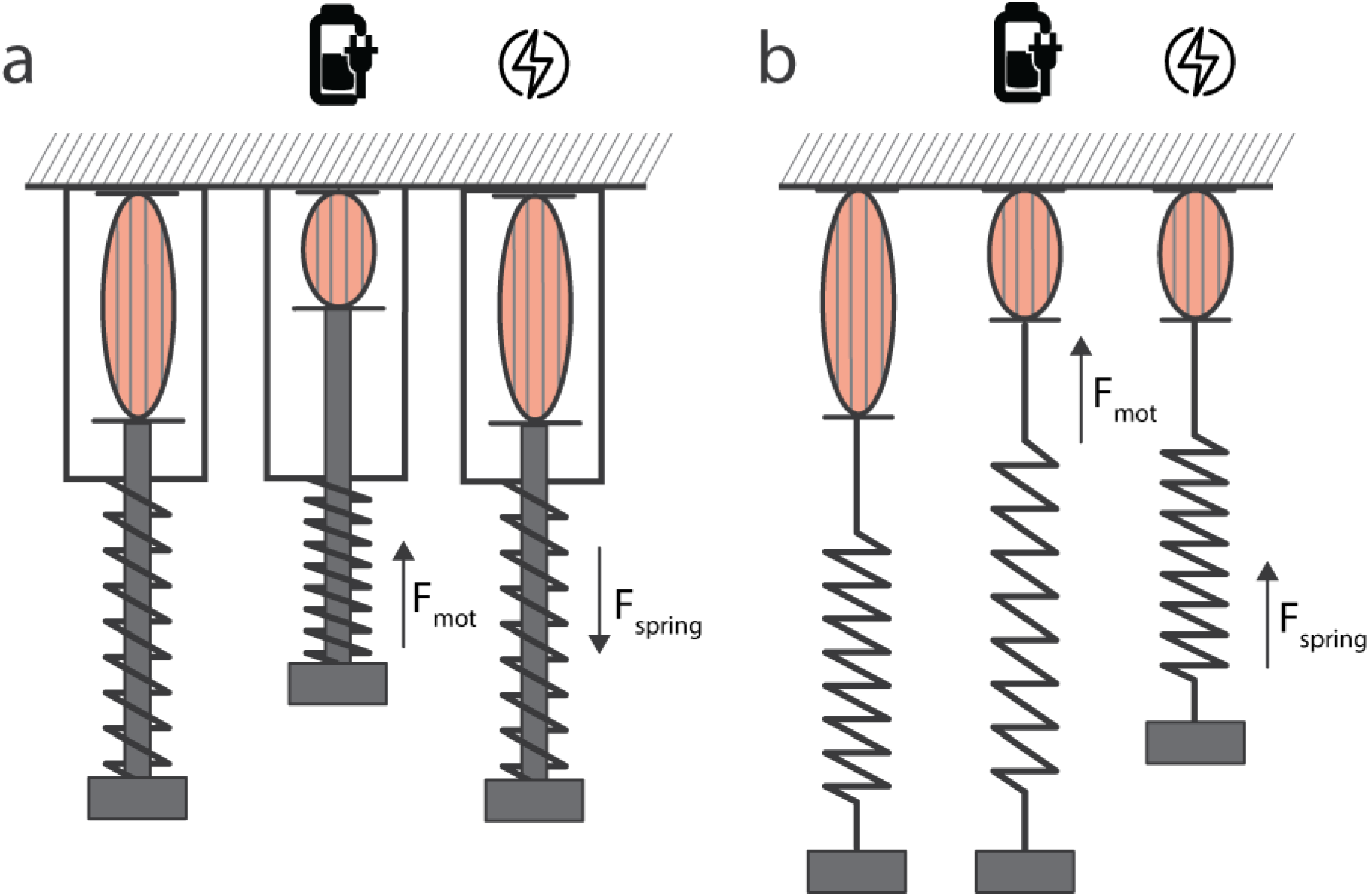
Parallel elastic and series elastic mechanisms for power modulation in comparison. (based on Yesilevskiy et al. 2015) a) Parallel elastic mechanism b) Series elastic mechanism. From left to right: relaxed state, preloading (energy is invested by the motor and stored in the spring), recovering (energy is returned by the spring to move the projectile). Note that in the parallel elastic mechanism, the position of the projectile is linked to the position of the motor. This is not the case in the series elastic mechanism, where spring and motor position are independent.

Both the predatory strike and the escape reaction of prey items showed a significant decrease in velocity (Figure 3,4) when temperature decreased. The average swimming velocity decreased by 79% in Ephemeroptera and by 45% in Zygoptera (as a prey item). Data on prey items are comparable to the performance decrease for the muscle-driven preparation phase of the predatory strike (50% in *A. imperator*, 71% in *E. najas*), but the protraction velocity only decreases by 34% in *A. imperator* and by 61 % in *E. najas,* respectively. Considering the data on ephemeropteran prey, odonatan predators would be at an advantage at lower temperatures even though the performance of the catapult mechanism deteriorates. Therefore, one would expect that the success rate of prey capturing increases with decreasing temperature. However, a study measuring the direct success rate of the predatory strike of Odonata larvae (with *Daphnia magna* as prey item) has shown that the capturing success does not depend on temperature (Quenta Herrera et al., 2018). Another study, focusing on the predation of larvae of the dragonfly *Hemicordulia tau* on the tadpoles of three frog species, showed a significant increase in predation with temperature, but only in the case of one prey species (Richards and Bull, 1990). The impact of predator-prey relationships has also been analysed in other arthropods (Kruse et al., 2008): while in carabids, the predation rate on *Drosophila* decreases with an increasing temperature, it increases in spiders (Kruse et al., 2008). In the predator-prey relationship between bass and mosquitofish, the predation rate decreases with an increase in temperature (Grigaltchik et al., 2012). These studies show that temperature-dependent predation rates not only differ in different predators, but also differ in the same predator with different prey items (Richards and Bull, 1990), highlighting the complexity of the interaction. We also found differences in the temperature dependence of mobility in different prey items (Figure 4). This might also lead to a differently composed prey spectrum of Odonata larvae at different temperature ranges (and therefore different times of the year), which has, to the best of our knowledge, not yet been analysed. Temperature influences many aspects of animal physiology (Huey and Kingsolver, 1989) in general, but here we clearly demonstrate that low temperature could mediate prey capturing in dragonfly larvae not only in the moment of the strike itself but also during stalking and targeting prey. The kinematic advantage of using a catapult driven prey capturing mechanism might keep the success rate of Odonata larvae constant at various temperatures and ensure continuous feeding during the year at various ambient temperatures. To fully understand the ecological implications of this mechanism, further field studies focusing on the prey spectrum of larvae in different Odonata species throughout the year are necessary.

## Conclusions

In the prehensile labial mask, a temperature-dependent catapult mechanism has been observed. This effect indicates a series elastic catapult which is not completely independent of muscle action. The mechanism is more similar to jumping motions in anurans than to other described parallel elastic catapult mechanisms in invertebrates, such as those of fleas or mantis shrimp, which are therefore more likely to be temperature independent. Slower trigger action and hydrodynamic forces might also be involved in the effect, but the effect of latter seems to be negligible. The decrease in performance is lower than in directly muscle-powered movements, such as the preparation of the strike and the avoidance reaction of prey items. This study sheds further light on the mechanics of prey capturing in Odonata larvae, as well as their ecological implications.

## Supporting information

Suppementary File 1

## Acknowledgements

We are grateful for the support by the members of the Functional Morphology and Biomechanics Group at Kiel University, especially Thies Büscher and Dennis Petersen. Thanks to Michelle Garitz for undergraduate research assistance and especially Lars Heepe for many elucidating discussions. Many thanks to Sophie Bodenstein, Helmut Jeske, Kai Lehmann and Arne Georg for providing specimens.

## Author Contributions

AK, LH, SNG and SB designed the study. AK, HLT and SB conducted the hs-video experiments. AK, HLT and SB carried out the analysis of the raw data. AK and LH carried out the statistical evaluation. AK, SNG and SB wrote the manuscript. All authors read, revised and approved the manuscript.

## Data Accessibility

All necessary data is available in the supplement. Raw datasets are available upon reasonable request from the corresponding author.

## Ethical Statement

No permission from research ethics or animal ethics committees was necessary. All applicable regulations concerning the protection of free-living species were followed.

## References

Astley, H.C., Roberts, T.J., 2012. Evidence for a vertebrate catapult: Elastic energy storage in the plantaris tendon during frog jumping. Biol. Lett. 8, 386–389. https://doi.org/10.1098/rsbl.2011.0982

Büsse, S., Hörnschemeyer, T., Gorb, S.N., 2017. The head morphology of Pyrrhosoma nymphula larvae (Odonata: Zygoptera) focusing on functional aspects of the mouthparts. Front. Zool. 14, 1–13. https://doi.org/10.1186/s12983-017-0209-x

Büsse, S., Koehnsen, A., Rajabi, H., Gorb, S.N., 2020. Hunting with catapults: the predatory strike of the dragonfly larva. bioRxiv 2020.05.11.087882. https://doi.org/10.1101/2020.05.11.087882

Corbet, P.S., 1999. Dragonflies: behaviour and ecology of Odonata. Harley books, Colchester

Dell, A.I., Pawar, S., Savage, V.M., 2014. Temperature dependence of trophic interactions are driven by asymmetry of species responses and foraging strategy. J. Anim. Ecol. 83, 70–84. https://doi.org/10.1111/1365-2656.12081

Galantis, A., Woledge, R.C., 2003. The theoretical limits to the power output of a muscle-tendon complex with inertial and gravitational loads. Proc. R. Soc. B Biol. Sci. 270, 1493–1498. https://doi.org/10.1098/rspb.2003.2403

Grigaltchik, V.S., Ward, A.J.W., Seebacher, F., 2012. Thermal acclimation of interactions: Differential responses to temperature change alter predator-prey relationship. Proc. R. Soc. B Biol. Sci. 279, 4058–4064. https://doi.org/10.1098/rspb.2012.1277

Gronenberg, W., 1996. Fast actions in small animals: springs and click mechanisms. J. Comp. Physiol. A 178, 727–734. https://doi.org/10.1007/BF00225821

Haldane, D.W., Plecnik, M., Yim, J.K., Fearing, R.S., 2016. A power modulating leg mechanism for monopedal hopping. IEEE Int. Conf. Intell. Robot. Syst. 2016-Novem, 4757–4764. https://doi.org/10.1109/IROS.2016.7759699

Hirano, B.Y.M., Rome, L.C., 1984. Jumping performance of frogs (*Rana pipiens*) as a function of muscle temperature. J. Exp. Biol. 108, 429–439.

Hoerner, S.F., 1965. Fluid-Dynamic Drag. Published by the author, Midland Park, NJ.

Huey, R.B., Kingsolver, J.G., 1989. Evolution of thermal sensitivity of ectotherm performance. Trends Ecol. Evol. 4, 131–135. https://doi.org/10.1016/0169-5347(89)90211-5

Ilton, M., Bhamla, M.S., Ma, X., Cox, S.M., Fitchett, L.L., Kim, Y., Koh, J., Krishnamurthy, D., Kuo, C., Temel, F.Z., Crosby, A.J., Prakash, M., Sutton, G.P., Wood, R.J., Azizi, E., Bergbreiter, S., Patek, S.N., 2018. The principles of cascading power limits in small, fast biological and engineered systems. Science 80, 1082. https://doi.org/10.1126/science.aao1082

Jara, F.G., Perotti, M.G., 2010. Risk of predation and behavioural response in three anuran species: Influence of tadpole size and predator type. Hydrobiologia 644, 313–324. https://doi.org/10.1007/s10750-010-0196-9

Josephson, R.K., 1993. Contraction dynamics and power output of skeletal-muscle. Annu. Rev. Physiol. 55, 527–546.

Koehnsen, A., Kambach, J., Büsse, S., 2020. Step by step and frame by frame - workflow for efficient motion tracking of high-speed movements in animals. Zoology 141, 125800 https://doi.org/10.1016/j.zool.2020.125800

Kovač, M., Schlegel, M., Zufferey, J.C., Floreano, D., 2010. Steerable miniature jumping robot. Auton. Robots 28, 295–306. https://doi.org/10.1007/s10514-009-9173-4

Kruse, P.D., Toft, S., Sunderland, K.D., 2008. Temperature and prey capture: Opposite relationships in two predator taxa. Ecol. Entomol. 33, 305–312. https://doi.org/10.1111/j.1365-2311.2007.00978.x

Pritchard, G., 1965. Prey capture by dragonfly larvae (Odonata; Anisoptera). Can. J. Zool. 43.2, 271–289

Pritchard, G., 1964. The prey of dragonfly larvae (Odonata; Anisoptera) in ponds in Northern Alberta. Can. J. Zool. 42, 785–800. https://doi.org/10.1139/z64-076

Quenta Herrera, E., Casas, J., Dangles, O., Pincebourde, S., 2018. Temperature effects on ballistic prey capture by a dragonfly larva. Ecol. Evol. 8.8. 4303–4311 https://doi.org/10.1002/ece3.3975VOLUME?

Rall, B.C., Brose, U., Hartvig, M., Kalinkat, G., Schwarzmüller, F., Vucic-Pestic, O., Petchey, O.L., 2012. Universal temperature and body-mass scaling of feeding rates. Philos. Trans. R. Soc. B Biol. Sci. 367, 2923–2934. https://doi.org/10.1098/rstb.2012.0242

Rall, B.C., Vucic-Pestic, O., Ehnes, R.B., Emmerson, M., Brose, U., 2009. Temperature, predator-prey interaction strength and population stability. Glob. Chang. Biol. 16, 2145–2157. https://doi.org/10.1111/j.1365-2486.2009.02124.x

Renaud, J.M., Stevens, E.D., 1983. The extent of long-term temperature compensation for jumping distance in the frog, Rana pipiens, and the toad, *Bufo americanus*. Can. J. Zool. 61, 1284–1287. https://doi.org/10.1139/z83-172

Richards, S.J., Bull, C.M., 1990. Size-limited predation on tadpoles of three australian frogs. Copeia 1990, 1041–1046. https://doi.org/10.2307/1446487

Rumble, J.R., Lide, D.R., Bruno, T.J., 2017. CRC handbook of chemistry and physics, 97th ed. CRC Press, New York.

Snodgrass, R.E., 1955. The dragonfly larva. Smithson. Misc. Collect. Washingt. D.C.

Sutton, G.P., Burrows, M., 2011. Biomechanics of jumping in the flea. J. Exp. Biol. 214, 836–847. https://doi.org/10.1242/jeb.052399

Tanaka, Y., Hisada, M., 1980. The hydraulic mechanism of the predatory strike in dragonfly larvae. J. Exp. Biol. 88, 1–19.

Vogel, S., 1997. Life in moving fluids, 2nd Edition. Princeton University Press, Princeton.

Vucic-Pestic, O., Ehnes, R.B., Rall, B.C., Brose, U., 2011. Warming up the system: Higher predator feeding rates but lower energetic efficiencies. Glob. Chang. Biol. 17, 1301–1310. https://doi.org/10.1111/j.1365-2486.2010.02329.x

Wiggins, G., Mackay, R., Smith, I., 1980. Evolutionary and ecological strategies of animals in annual temporary pools. Arch. für Hydrobiol. Suppl. 58.1, 97–206

Yesilevskiy, Y., Xi, W., Remy, C.D., 2015. A comparison of series and parallel elasticity in a monoped hopper. Proc. - IEEE Int. Conf. Robot. Autom. 2015-June, 1036–1041. https://doi.org/10.1109/ICRA.2015.7139304

