## Supplementary material for "The temperature-dependent predatory strike of Odonata larvae": Suppementary File 1

**Supplementary material S1**

**Drag Force Estimate**

**1.1 Impact of density:**

According to the drag equation, the drag force $F_d$ can be calculated as follows:

$$F_d = \frac{1}{2}\,\rho v^2 C_d A$$

$F_d$ depends on the density ($\rho$) the flow velocity ($v$), the specific area $A$ and the drag coefficient $C_d$. $\rho$ is the only variable that is a direct function of temperature. To calculate the increase in drag due to lower temperatures ($\Delta F_d$), the drag equation can be simplified to the following:

$$\Delta F_d = \Delta\rho * \mathrm{const.}$$

In the following, we look at an exemplary change from 10°C to 20°C, similar to the range in the experiments. (Data from: Rumble et al. 2017. CRC handbook of chemistry and physics, 97th edition)

| | Temperature [°C] | Density [g/ml] |
|---|---|---|
| | 20 | 0.998207 |
| | 10 | 0.999702 |
| Δ | - 10 | 0.001495 |

This relates to a relative increase of 0.001498, or 0.15%. As $\rho$ is incorporated linearly into the drag equation, decreasing the temperature by ten degrees relates to a neglectable increase of 0.15% in drag.

**1.2 Impact of Re change:**

The drag coefficient $C_d$ is an object specific constant, which has to be estimated via flow channel or CFD analyses. It is not temperature dependent per se, but it is a function of the Reynolds number (Re) which is dependent on temperature. To roughly estimate the impact of a temperature change of ten degrees, we calculated an object with the approximate maximum width of the prehensile labial mask of both *Anax* (7.5mm) and *Erythromma* (1.6mm) to assess how much Re changes might change $C_d$.

The Reynolds Number is calculated as follows:

$$Re = \frac{\rho\, \upsilon\, d}{\eta}$$

with the characteristic length ($d$). The velocity ($\upsilon$) was determined from our velocity data.

Dynamic viscosity ($\eta$) and density ($\rho$) were taken from literature data of freshwater at 25°C.

| | T [°C] | l [mm] | ρ [g/ml] | υ [m/s] | η [mPa*s] | *Re* |
|---|---|---|---|---|---|---|
| *Erythromma* | 20 | 1.6 | 0.999702 | 0.4 | 1.0016 | ~ 650 |
| | 10 | 1.6 | 0.998207 | 0.4 | 1.3059 | ~ 500 |
| *Anax* | 20 | 7.5 | 0.999702 | 0.3 | 1.0016 | ~ 2200 |
| | 10 | 7.5 | 0.998207 | 0.3 | 1.3059 | ~ 1700 |

*Re* decreases by around 23% in *Erythromma* as well as in *Anax* with a temperature decrease of 10°C.
