## Supplementary Figure 2 for "The temperature-dependent predatory strike of Odonata larvae"

**Supplementary material S2**

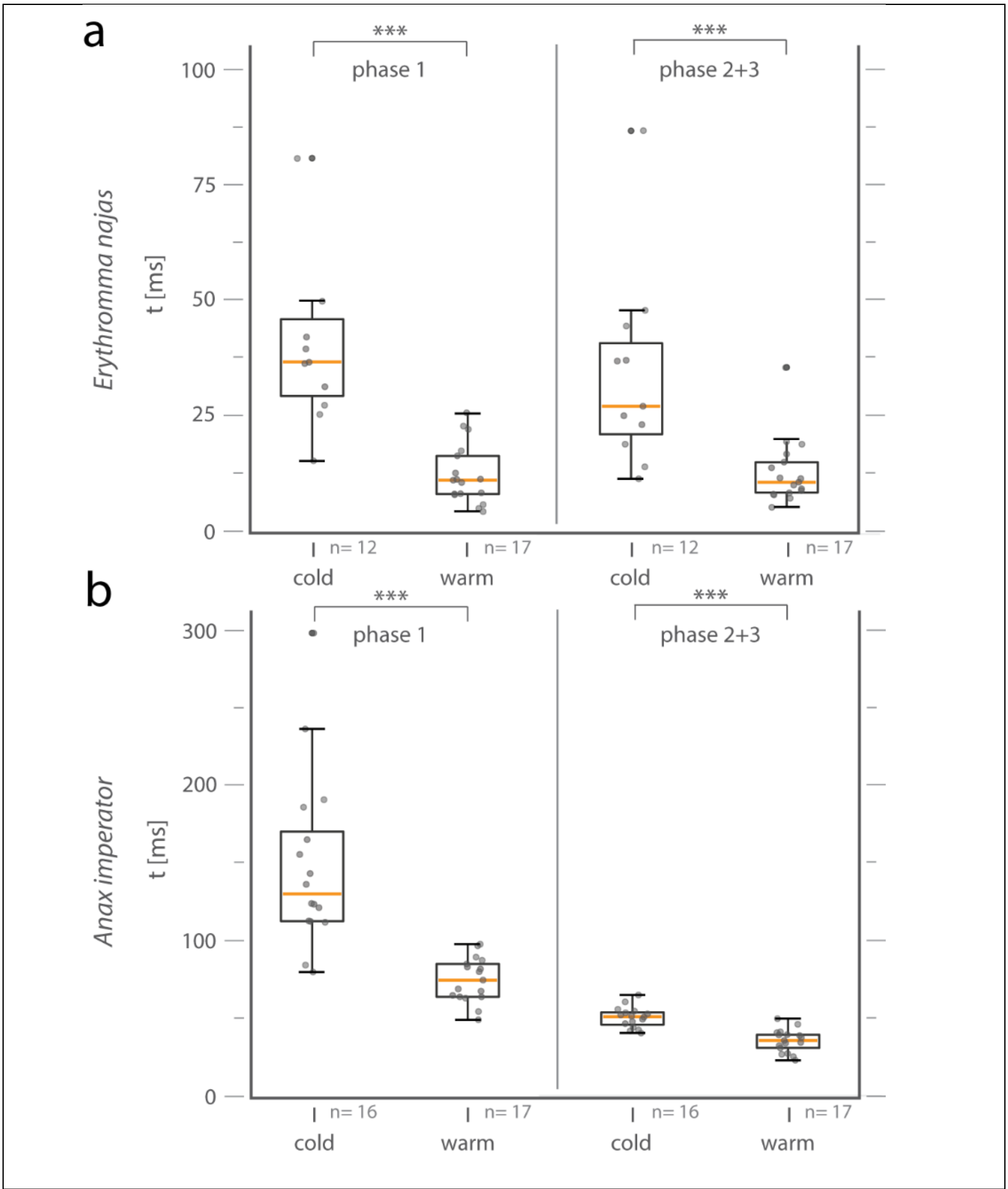


**S2: Supplementary figure: Phase duration increases with a decrease in ambient temperature.** Duration of phase 1 (opening of the labial palps) versus phase 2+3 (full extension of the prehensile labial mask). For both strike phases, the duration decreases significantly with an increase in temperature. a) *E. najas*: phase 1: Mann-Whitney Rank Sum Test, $p<0.001$ n=17/12; phase 2+3: Mann-Whitney Rank Sum Test, $p<0.001$ n=17/12). b) *A. imperator*: phase 1: Mann-Whitney Rank Sum Test, $p<0.001$ n=17/16; phase 2+3: t-test, $p<0.001$ n=17/16), box ends define 25$^{th}$ and 75$^{th}$

percentiles, with a line highlighting the median and error bars at the 10th and 90th percentiles. Dots indicate individual data points.
